# Eating junk-food has opposite effects on intrinsic excitability of nucleus accumbens core medium spiny neurons in obesity-susceptible vs. -resistant rats

**DOI:** 10.1101/658971

**Authors:** Max F. Oginsky, Carrie R. Ferrario

## Abstract

The nucleus accumbens (NAc) plays critical roles in motivated behaviors, including food-seeking and feeding. Differences in NAc function contribute to over-eating that drives obesity, but the underlying mechanisms are poorly understood. In addition, there is a fair degree of variation in individual susceptibility vs. resistance to obesity that is due in part to differences in NAc function. For example, using selectively bred obesity-prone and obesity-resistant rats, we have found that excitability of medium spiny neurons within the NAc core is enhanced in obesity-prone vs. resistant populations, prior to any diet manipulation. However, it is unknown whether consumption of sugary, fatty junk-food alters MSN excitability. Here, whole-cell patch clamp recordings were conducted to examine MSN intrinsic excitability in adult male obesity-prone and obesity-resistant rats with and without exposure to a sugary, fatty junk-food diet. We replicated our initial finding that basal excitability is enhanced in obesity-prone vs. obesity-resistant rats and determined that this is due to a lower I_*A*_ in prone vs. resistant groups. In addition, the junk-food diet had opposite effects on excitability in obesity-prone vs. obesity-resistant rats. Specifically, junk-food enhanced excitability in MSNs of obesity-resistant rats; this was mediated by a reduction in I_*A*_. In contrast, junk-food reduced excitability in MSNs from obesity-prone rats; this was mediated by an increase in I_*KIR*_. Thus, individual differences in obesity-susceptibility influence both basal excitability and how MSN excitability adapts to junk-food consumption.

## I. INTRODUCTION

The nucleus accumbens (NAc) mediates motivational responses to rewards such as food, as well as to reward-associated stimuli. This is due in part of the ability of medium spiny neruons (MSNs) within this region to integrate glutamatergic and dopaminergic signals from mesolimbic and cortical inputs. However, another factor that influences this integration, and ultimately NAc output, is intrinsic excitability of MSNs. Within MSNs, excitability is shaped largely by inwardly rectifying potassium currents (I_*KIR*_), which help maintain their hyperpolarized states, and the fast transient potassium current (I_*A*_), which regulates the time to the first action potential following depolarization (John and Manchanda 2011; Nisenbaum and Wilson 1995). Dopaminergic transmission bi-directionally influences MSN excitability via indirect actions on I_*KIR*_ and I_*A*_ (Azdad et al. 2009; Perez et al. 2006; Podda et al. 2010). For example, activation of D1- or D2-type dopamine receptors (D1R, D2R) can inhibit I_*KIR*_, resulting in increased excitability (Perez et al. 2006; Podda et al. 2010), while activation of D2Rs can also enhance I_*A*_ to reduce excitability (Perez et al. 2006). Furthermore, repeated exposure to cocaine, which elevates dopamine levels in the NAc, reduces MSN intrinsic excitability by modulating a number of ionic conductances, including I_*KIR*_ and I_*A*_ (Henry and White 1992; Hu et al. 2004; Kourrich and Thomas 2009; Mu et al. 2010). However, less well understood is how consumption of sugary, fatty foods, which also enhance NAc dopamine (Bassareo et al. 2015; McCutcheon and Roitman 2019) and drive over-eating and obesity (Small 2009; Tomasi and Volkow 2013; Vainik et al. 2013), may influence MSN intrinsic excitability. This is particularly important given the role of the NAc in food-seeking and craving, which are enhanced following consumption of palatable foods (Derman and Ferrario 2018a; b).

Individual susceptibility to obesity has been linked to basal and diet-induced alterations in NAc function and enhanced cue-triggered food-seeking behaviors (Alonso-Caraballo et al. 2018; Ferrario 2017; Stice et al. 2008; Yokum et al. 2011). For example, motivational responses to food cues are stronger in rodent models of obesity susceptibility (Derman and Ferrario 2018a; Robinson et al. 2015). This is associated with enhanced responsivity of mesolimbic circuits (Oginsky et al. 2016b; Vollbrecht et al. 2015), and greater intrinsic excitability of MSNs in the NAc of obesity-prone vs. obesity-resistant groups prior to diet manipulation (Oginsky et al. 2016b). These data are consistent with stronger striatal activations in response to food-cues in overweight and obesity-susceptible human populations (Demos et al. 2012; Vainik et al. 2013). However, how consumption of sugary, fatty foods may affect excitability of MSNs in models that capture individual susceptibility to obesity is unknown.

In the current study, we determined how excitability of MSNs in the NAc core of male obesity-prone and obesity-resistant rats is altered by brief access to a sugary, fatty, junk-food diet. This diet was chosen because it produces enhancements in NAc glutamatergic transmission in obesity-prone but not obesity resistant rats (Oginsky et al. 2016a) that are similar to those seen following repeated cocaine exposure (Alonso-Caraballo et al. 2018; Ferrario 2017).

Consistent with previous results (Oginsky et al. 2016b), we found that MSNs were hyper-excitable in obesity-prone compared to obesity-resistant rats maintained on standard lab chow; this was due in part to lower I_*A*_ in obesity-prone vs. obesity-resistant rats. Consumption of the junk-food diet for just 10 days resulted in reduced excitability in obesity-prone rats, but enhanced excitability in obesity-resistant rats compared to their chow fed counterparts. Furthermore, junk-food-induced increases in intrinsic excitability in obesity-resistant rats were due to a decrease in I_*A*_, whereas diet-induced reductions in excitability in obesity-prone rats were due to an increase in I_*KIR*_. These data are discussed in light of the role of the NAc in motivation and interactions between individual susceptibility to obesity and diet-induced plasticity in this region.

## II. MATERIALS AND METHODS

Subjects: Rats were pair-housed, maintained on a reverse light-dark schedule (12/12), and all tests were conducted during the dark phase starting 3-4 h after dark onset. Obesity-prone and obesity-resistant rats were bred inhouse by the University of Michigan breeding core. These lines were originally established by Barry Levin (Levin et al. 1997), and obesity-prone and obesity-resistant phenotypes have been confirmed by our group (Alonso-Caraballo et al. 2018; Oginsky et al. 2016b; Vollbrecht et al. 2015). Rats had free access to food and water throughout. All procedures were approved by The University of Michigan Committee on the Use and Care of Animals. All drugs and reagents were obtained from Sigma (St. Louis, MO, USA) or Tocris (Minneapolis, MN, USA) unless otherwise stated.

Whole-cell patch clamp recordings of MSNs were made from adult (*P*70-*P*80) males given free access to standard lab chow (certified rodent diet 5001; Lab Diet, St. Louis, MO; 200g; % of calories: 19.6% fat, 14% protein, 58% carbohydrates; 4.5kcal/g) or a junk-food diet made in house. The junk-food diet consisted of Ruffles original potato chips (40g), Chips Ahoy original chocolate chip cookies (130g), Jif smooth peanut butter (130g), Nesquik powdered chocolate flavoring (130g), powdered Lab Diet 5001 (200g; % of calories: 19.6% fat, 14% protein, 58% carbohydrates; 4.5kcal/g), and water (180ml) combined in a food processor. Rats were either given ad libitum access to junk-food for 10 days, or maintained on standard lab chow. During this time, food intake per cage and body weight were measured daily.

All food was removed from the home cage 14-16 hrs prior to slice preparation in order to avoid potential confounds of differences in food consumption on the recording day. Recordings were conducted in the presence of the GABA receptor antagonist picrotoxin (50*μM*) as previously described (Oginsky et al. 2016b). Briefly, rats were anesthetized with chloral hydrate (400 mg/kg, i.p.), the brains were rapidly removed and placed in ice cold oxygenated (95% O2 - 5% CO2) aCSF containing (in mM) 125 NaCl, 25 NaHCO3, 12.5 glucose, 1.25 NaH2PO4, 3.5 KCl, 1 L-ascorbic acid, 0.5 CaCl2, 3 MgCl2, and 305 mOsm; pH 7.4. Coronal slices (300 microns) containing the NAc were made using a Leica VT1200 vibratory microtome (Leica Biosystems, Buffalo Grove, IL, USA) and allowed to rest in oxygenated aCSF for at least 40 min before recording. For the recording aCSF (2 ml/min), CaCl2 was increased to 2.5 mM and MgCl2 was decreased to 1 mM. Patch pipettes were pulled from 1.5-mm borosilicate glass capillaries (WPI, Sarasota, FL) to a resistance of 3-7 MΩ with a horizontal puller (model P97, Sutter Instruments, Novato, CA, USA) and filled with a solution containing (in mM) 130 K gluconate, 10 KCl, 1 EGTA, 2 Mg2+-ATP, 0.6 Na+ -GTP, and 10 HEPES, pH 7.3, 285 mOsm for current clamp recordings and for voltage clamp recordings.

For current clamp studies, MSNs in the NAc core were identified based on resting membrane potential and action potential firing in response to current injection (200 to 175 or 275 pA, 25 pA increments, 1000 ms (Nisenbaum and Wilson 1995; Wilson and Kawaguchi 1996). Cell health was evaluated by resting membrane potential, action potential amplitude, action potential width, input resistance. Any recordings where these changed during the course of the experiment were rejected from further analysis. Measurements were taken from the first action potential elicited by the minimum current injection or between the first two action potentials for interspike interval and afterhyperpolarization (AHP).

The I/V relationship was determined by calculating the difference between the baseline voltage and the voltage at 200 ms after current injection between −200 mV and +140 mV. Neuronal excitability was determined by measuring the number of action potentials elicited by each depolarizing current injection. Input resistance was determined by the change in voltage from −50 to +50 pA current injections. Rheobase is defined as the minimum amount of current injection to elicit an action potential. The action potential threshold was determined using the maximum second derivative method (Sekerli et al. 2004). The peak amplitude was defined as the difference between the action potential peak and threshold. The interspike interval was measured between the peak amplitude of the first two action potentials in a train of the first current injection that elicited spikes. The amplitude of the AHP was measured from the action potential threshold to the lowest potential of the AHP.

For voltage clamp studies, I_*KIR*_ was recorded using a ramp voltage protocol (−150 to −50 mV) at a rate of 100 mV/sec starting from a holding potential of −70 mV before and after 1 mM BaCl2 administration. The recording after BaCl2 was subtracted from the baseline recording to obtain I_*KIR*_ (Jin et al. 2013; Mermelstein et al. 1998). To measure I_*A*_, the voltage dependence of activation was measured by a series of sweeps in which a hyperpolarizing prepulse to −120 mV for 100 ms was followed by a 1000 ms depolarizing test pulse that ranged from −50 to +60 mV with 10 mV increments. Inactivation was measured with a series of sweeps in which the membrane potential was stepped to a prepulse between −120 and +20 mV with 10 mV increments (100 ms) followed by a constant test pulse to +20 mV (1000 ms). I_*A*_ was isolated by including 20 mM tetraethylammonium (TEA) to block delayed-rectifying potassium currents, 1 uM tetrodotoxin (TTX) to block voltage-gated sodium currents. In addition, calcium was reduced from 2 mM to 0.5 mM in the recording aCSF to minimize calcium currents. I_*A*_ was further isolated by digital subtraction of the leak conductance. To measure activation, I_*A*_ was evoked by test pulses between +50 and −60 mV from +50 mV holding potential without a negative prepulse. These current traces were subtracted from those evoked with negative prepulse. To measure inactivation, the subtraction protocol contained a depolarizing prepulse to +20 mV before the test pulse to −20 mV. The conductance across voltages for activation and inactivation were fitted with a first order Boltzmann equation. We did not correct for liquid junction potential. Analyses were made from single sweeps. Ns for electrophysiological data are given as the number of rats followed by the number of cells (e.g., N=6,8 indicates 8 cells from 6 rats).

## III. RESULTS

Obesity-resistant (OR) and obesity-prone (OP) rats were given ad libitum access to either junk-food (OR N=8, OP N=9) or chow (OR N=7, OP N=9) for 10 days prior to slice preparation. As expected, rats in the obesity-prone group consumed more food compared to obesity-resistant rats (Fig. 1A: Two-way ANOVA; main effect of line: *F*_(1,29)_=24.1, p<0.001) and rats with access to junk-food consumed more food than rats in the chow groups (Fig. 1A: Two-way ANOVA; main effect of condition, *F*_(1,29)_=45, p<0.001). In addition, differences between obesity-prone and obesity-resistant groups were primarily driven by significantly greater junk-food consumption in obesity-prone vs. obesity-resistant rats (Sidak’s post-test; OP JF vs OR JF p=0.003). Total weight gained across the 10 day feeding period is shown in Fig 1B. Junk-food groups gained more weight than their chow-fed counterparts (Fig. 1B: Two-way ANOVA; main effect of diet, *F*_(1,29)_=40.5, p<0.0001) and obesity-prone rats gained significantly more weight than obesity-resistant groups (Fig. 1B: Two-way ANOVA; main effect of line, *F*_(1,29)_=11, p<0.01).

**FIG. 1:**
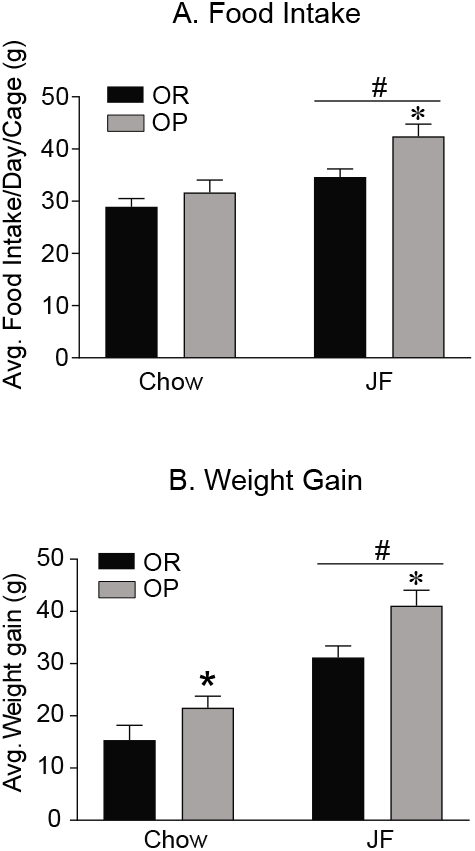
Food intake and weight gain in obesity-resistant (OR) and obesity-prone (OP) rats. A) Estimated average daily food intake in OR and OP fed either chow or junk-food (JF). Rats given junk-food eat more than their chow counterparts, and rats in the OP group consume more junk-food than those in the OR group. B) Average weight gain across all groups. As expected, OP rats are heavier than ORs, and weight gain is greater in junk-food vs. chow fed groups. OP chow = 9 rats, OP JF = 9 rats, OR Chow = 7 rats, OR JF = 8 rats, = main effect of junk-food, p <0.05; *=OP vs OR p<0.05; all data shown as average (SEM), unless otherwise noted.

We next examined intrinsic excitability of MSNs from obesity-prone and obesity-resistant rats fed either chow or junk-food by measuring the number of action potentials elicited by increasing current injection, current/voltage relationships, and intrinsic membrane properties (OP-Chow N =4,10; OP-JF N=4,10; OR-Chow N=4,12; OR-JF N=5,12). Consistent with our previous results (Oginsky et al. 2016b), the number of action potentials elicited across current injections was significantly greater in MSNs from adult obesity-prone vs. obesity-resistant chow groups (Fig. 2A: Two-Way RM ANOVA; significant group x current injection interaction: *F*_(10,200)_=4.4, p<0.05; Sidak’s posttest, p<0.001). When comparisons were made between obesity-prone and obesity-resistant junk-food groups, this relationship was reversed, such that MSNs were more excitable in obesity-resistant vs. obesity-prone junk-food fed groups (Fig. 2B: Two-Way RM ANOVA; significant condition x current injection interaction: *F*_(10,200)_=4.9, p<0.001; Sidak’s post-test, p<0.001). This was due to opposite effects of junk-food in obesity-resistant and obesity-prone groups. Specifically, when we compared the number of action potentials elicited across current injections within obesity-resistant groups given chow or junk-food, we found a significant increase in excitability (Fig. 2C: Two-way RM ANOVA; significant condition x current injection interaction: *F*_(10,220)_=3.1; p=0.001; Sidak’s post-test, p<0.05). Conversely, in obesity-prone rats junk-food consumption resulted in a significant reduction in excitability (Fig. 2D: Two-way RM ANOVA; significant condition x current injection interaction: *F*_(10,220)_=5.9; p<0.001; Sidak’s posttest, p<0.001). Example traces from each group are shown in Fig. 2E and 2F. In sum, basal intrinsic excitability was enhanced in obesity-prone vs. obesity-resistant rats, and junk-food diet consumption produced opposite effects on excitability in these lines.

**FIG. 2:**
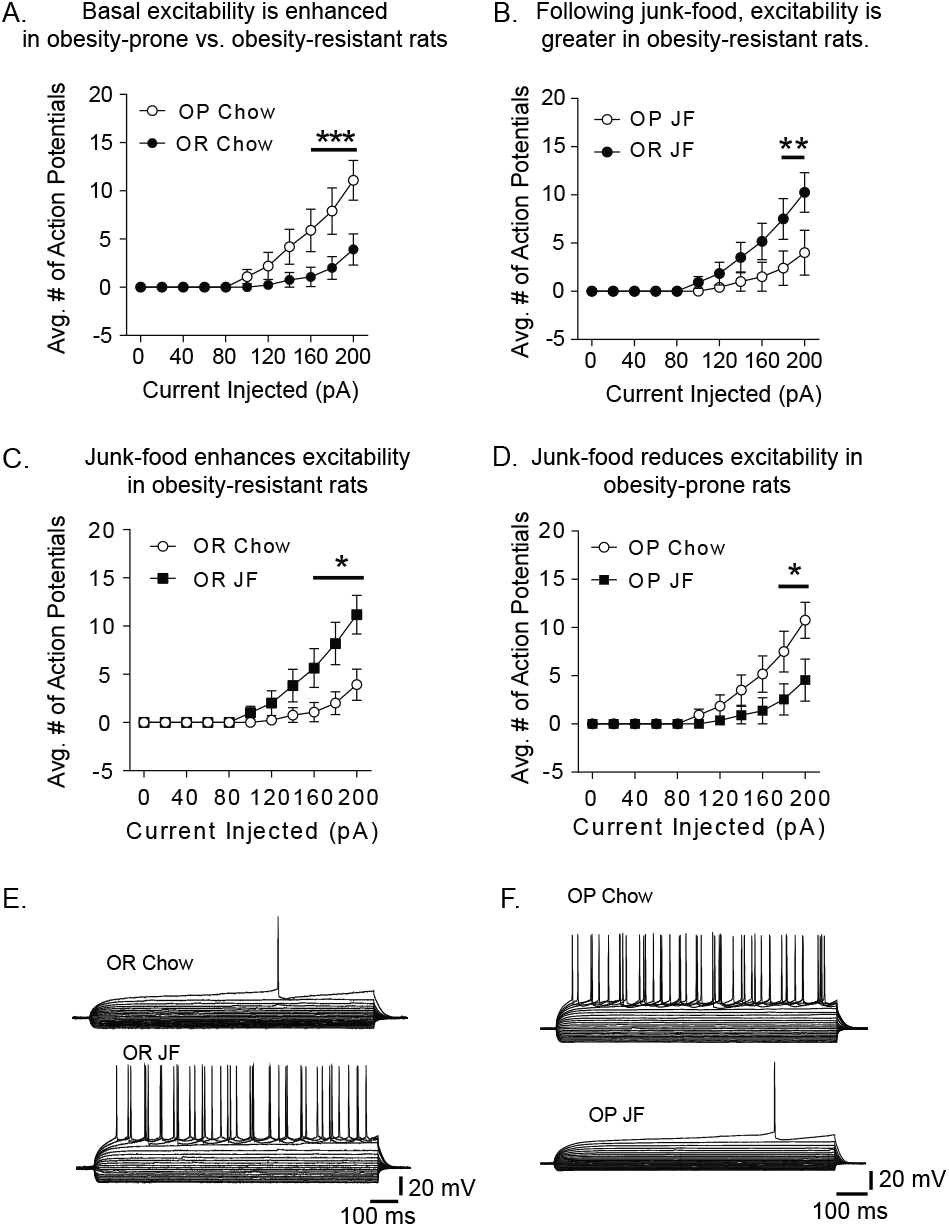
Junk-food (JF) increases MSN excitability in obesity-resistant (OR) rats whereas junk-food decreases MSN excitability in obesity-prone (OP) rats. A) The average number of action potentials per current injected (0 to 200 pA) in MSNs from chow fed obesity-resistant (OR) and obesity-prone (OP) rats. Basal excitability was greater OP vs OR chow fed groups. B) The average number of action potentials per current injected (0 to 200 pA) in MSNs of junk-food fed groups. Following junk-food consumption, excitability is lower in OP vs OR groups. C) Comparisons between OR chow and OR junk-food groups. Junk-food increases excitability in OR rats. D) Comparisons between OP chow and OP junk-food groups. Junk-food reduces excitability in OP rats. E) Example traces from OR chow and OR junk-food rats. F) Example from OP chow and OP junk-food rats. Number of cells and rats per groups. OP Chow = 4,10; OP JF = 4,10; OR Chow = 4,12; OR JF = 4,13. *=p<0.05, **=p<0.01,***=p<0.001.

To identify potential mechanisms underlying effects of junk-food on excitability, we next examined current-voltage relationships and intrinsic membrane properties within obesity-resistant (Fig. 3A) and obesity-prone lines (Fig. 3B). Consistent with an increase in the number of action potentials in obesity-resistant chow vs. junk-food groups, the I/V curve showed a greater change in membrane potential (Vm) in response to current injection in MSNs from obesity-resistant junk-food vs. chow groups (Fig. 3A: Two-way RM ANOVA; significant condition x current injection interaction: *F*_(17,391)_=9.8, p<0.0001). These differences were seen at both negative and positive current injections (Sidak’s post-test, p<0.0001). The opposite was seen when comparisons were made within obesity-prone groups, with smaller changes in membrane potential found in response to both positive and negative current injections between obesity-prone junk-food vs. chow groups (Fig. 3B: Two-way RM ANOVA; significant condition x current injection interaction: *F*_(17,306)_=7.4, p<0.0001; Sidak’s post-test, p<0.0001). Consistent with effects on excitability and I/V relationships, the rheobase (i.e., the minimal amount of current to elicit an action potential) was decreased in MSNs from obesity-resistant junk-food vs. chow groups, whereas in the obesity-prone groups, the rheobase was increased in junk-food compared to chow groups (Fig. 3C: Two-way ANOVA; significant line x diet interaction: *F*_(1,40)_ = 10.6, p<0.01; OR chow vs. OR JF, Sidak’s post-test, p ¡ 0.05; OP chow vs. OP JF, Sidak’s post-test, p<0.05). In addition, input resistance was significantly increased in obesity-resistant junk-food vs. chow groups (Fig. 3D, Two-way ANOVA; significant line x diet interaction: *F*_(1,40)_=8.0, p<0.01; OR Chow vs. OR JF: Sidak’s post-test, p<0.05), with trends towards a reduction in input resistance in obesity-prone junk-food vs. chow groups (OP Chow vs. OP JF: Sidak’s post-test, p=0.054). Finally, although junk-food did not alter the resting membrane potential (RMP) in obesity-resistant rats, the RMP was significantly hyperpolarized in MSNs from obesity-prone junk-food vs. obesity-prone chow groups (Fig. 3E: Two-way ANOVA; significant line x diet interaction: *F*_(1,40)_ =5.3, p<0.05; OP JF vs. OP chow, Sidak’s post-test, p<0.05). No group differences in action potential threshold were found (Fig. 3F.) Thus, effects of junk-food on excitability within obesity-resistant rats appear to be due to reductions in rheobase and increases in input resistance, whereas effects in obesity-prone rats appear to be due to increases in rheobase and a hyperpolarized resting membrane potential. Because there was a greater change in membrane potential at positive current injections and a decrease in rheobase in MSNs from obesity-resistant junk-food vs. obesity-resistant chow groups, we hypothesized I_*A*_ may be decreased following junk-food diet consumption. To test this, we performed voltage-clamp experiments to measure I_*A*_ (see methods for details). The I_*A*_ amplitude was significantly decreased in MSNs from obesity-resistant junk-food vs. chow groups (Fig. 4A: unpaired t-test, *t*_12_=2.4, p<0.05; OR Chow N=3,7; OR JF N=3,7). In addition, while the I_*A*_ inactivation curve was similar between chow and junk-food groups, the I_*A*_ activation curve was shifted to the right in MSNs from obesity-resistant junk-food vs. chow fed groups (Fig. 4B: Two-way RM ANOVA; significant main effect of diet: *F*_(1,6)_=11.6, p<0.05). This shift resulted in a smaller window current (see inset Figure 4B) in the junk-food group that is consistent with a significant reduction in the membrane potential for half-activation in these cells (Fig. 4C; unpaired t-test, *t*_12_=2.3; p<0.05). Example traces are shown in panel D. Taken together these data suggest that JF-diet induced increases in intrinsic excitability in obesity-resistant rats are mediated in part by a reduction in I_*A*_.

**FIG. 3:**
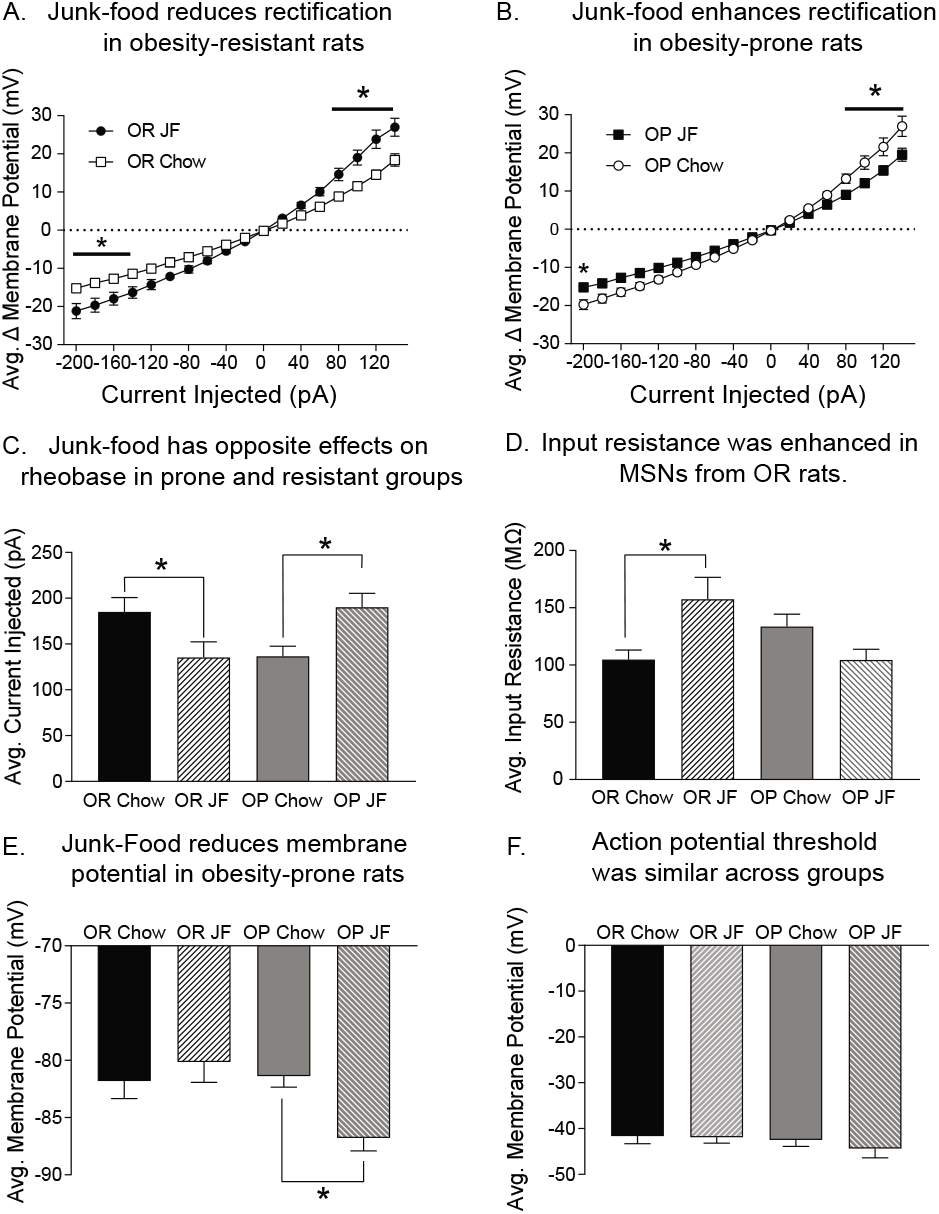
Junk-food (JF) produces opposite changes in intrinsic membrane properties in obesity-resistant (OR) and obesity-prone (OP) rats. A) The change in membrane potential at each current injected (−200 to +140 pA) in MSNs from OR rats fed either chow or junk-food. Junk-food increases rectification in OR rats. B) The change in membrane potential at each current injected (−200 to +140 pA) in MSNs from OP rats fed either chow or junk-food. Junk-food reduces rectification in OP rats. C) Average rheobase in chow and junk-food fed groups. Junk-food decreases rheobase in ORs, but increases it in OP rats. D) Average input resistance in chow and junk-food fed groups. Junk-food increased input resistance in OR rats and produced modest reductions in OP rats. E) Average resting membrane potential (RMP) in chow and junk-food fed groups. Resting membrane potential was decreased in OP rats fed junk-food vs. chow, but unaltered in ORs. F) Average action potential firing threshold was similar across groups. *=p<0.05.

**FIG. 4:**
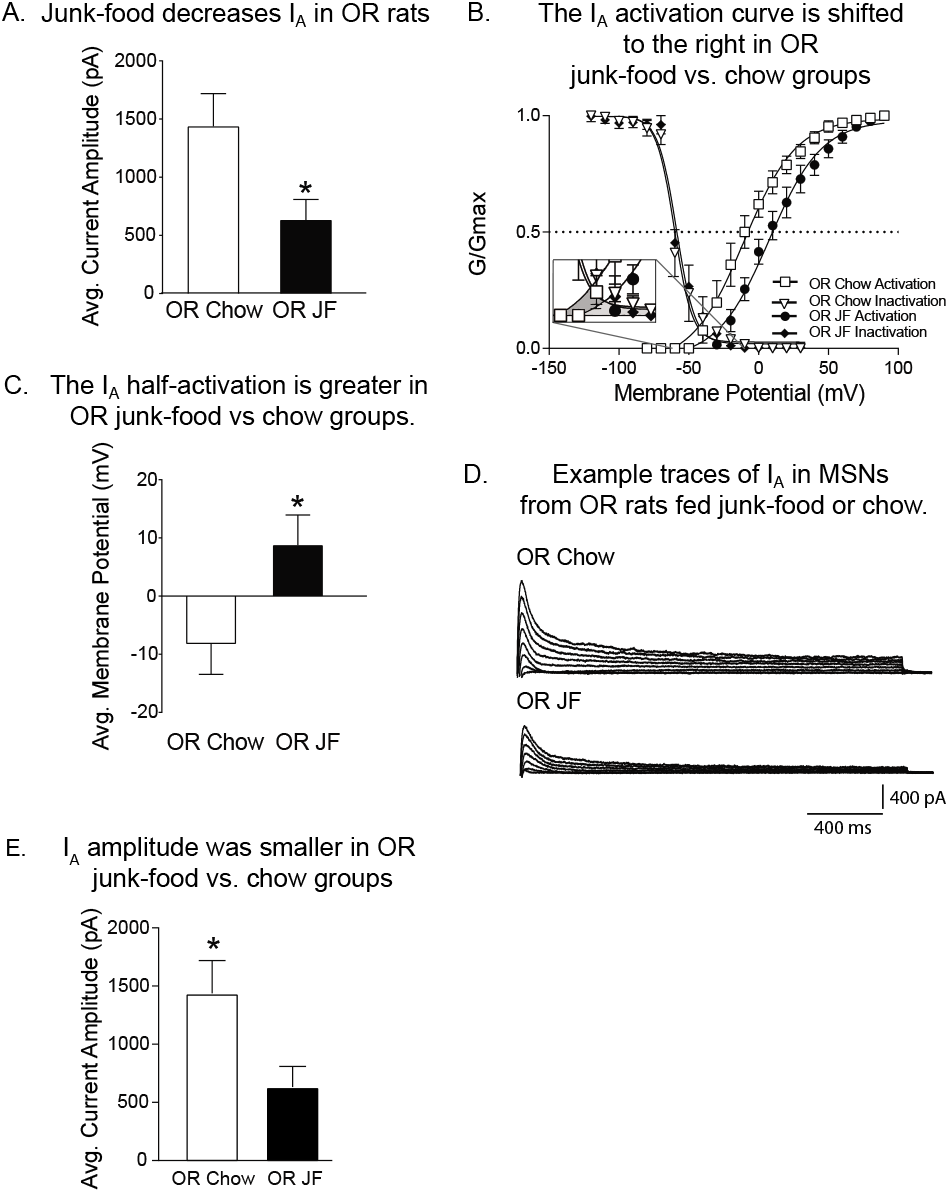
Junk-food reduces the A-current in MSNs from obesity-resistant (OR) rats. A) The average current magnitude average of I_*A*_ in obesity-resistant chow vs. junk-food groups. I_*A*_ current magnitude was decreased in MSNs from OR rats fed junk-food compared to OR rats fed chow. B) Normalized conductance-voltage plot of I_*A*_ activation and inactivation in OR junk-food and chow groups. Junk-food resulted in a shift to the right in the activation curve. C) The average half-activation in obesity-resistant chow vs. junk-food groups. The half-activation was ~15mV depolarized in MSNs from OR rats fed junk-food vs. chow. D) Example traces from MSNs in OR rats fed either junk-food or chow. E) Average I_*A*_ current amplitude. Junk-food decreased I_*A*_ current amplitude in MSNs from OR rats. Number of cells and rats per groups. OR Chow = 3,7; OR JF 4,7. *=p<0.05.

When the same analysis was conducted in MSNs from obesity-prone junk-food and chow groups, no differences in I_*A*_ amplitude (data not shown; 762.7 ±170 pA OP Chow vs. 758.8 ±148 pA OP JF; OP Chow N = 2,6; OP JF N = 3,6), or the half-activation voltages (6.9 ±4.2 mV OP Chow vs. 12.6 ±3.3 mV OP JF or half-inactivation voltages (−56.3 ±4.6 mV OP Chow vs. −58.8 ±0.48 mV OP JF) were found. This is surprising, given the effect the junk-food diet had on I_*A*_ in MSNs from obesity-resistant rats. However, in obesity-prone rats, junk-food diet resulted in a decrease in RMP of ~5 mV, and the RMP is predominantly regulated by I_*KIR*_ in MSNs (Uchimura et al. 1989; Uchimura and North 1990). Therefore, we hypothesized that increases in I_*KIR*_ may contribute to reduced excitability following junk-food consumption in obesity-prone rats. To examine this possibility, we used an established ramp protocol and pharmacological approaches to isolate I_*KIR*_ (Jin et al., 2013; Mermelstein et al., 1998; see methods for additional details). We found that I_*KIR*_ amplitude at −150 mV was enhanced in MSNs from obesity-prone rats fed either chow or junk-food (Fig. 5A: unpaired t-test, *t*_11_=4.05, p<0.01; OP Chow N=3,8; OP JF N=3,5). Figure 5B shows the average difference in I_*KIR*_ between chow and junk-food groups across the entire ramp protocol. This increase in I_*KIR*_ is consistent with junk-food induced reductions in excitability, RMP, and increased rheobase described above.

**FIG. 5:**
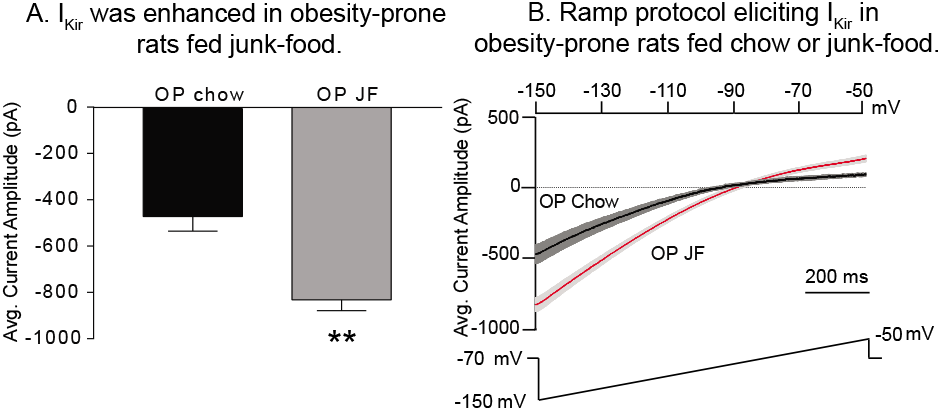
Junk-food diet enhances I_*KIR*_ in MSNs from OP rats. A) Average I_*KIR*_ amplitude in OP junk-food and chow groups. Consistent with reductions in excitability, junk-food enhanced I_*KIR*_ amplitude compared to the chow fed group. B) Average I_*KIR*_ amplitude in MSNs from OP junk-food (gray) and OP chow (Black) groups. The ramp protocol is shown below, with the membrane potential along the ramp shown above. Number of cells and rats per groups. OP Chow = 3,8; OP JF = 3,5. **=p<0.01.

## IV. DISCUSSION

MSNs in the NAc core are more excitable in adult male obesity-prone vs. obesity-resistant rats prior to any diet manipulation (Oginsky et al. 2016a). Here, we replicated this finding and determined that basal differences in excitability are due to lower I_*A*_ in obesity-prone vs. obesity-resistant groups. In addition, we examined how consumption of a junk-food diet affects MSN excitability in this model of individual susceptibility. We found that after the junk-food diet, there was enhanced excitability in MSNs of obesity-resistant rats; this was mediated by a reduction in I_*A*_. In contrast, junk-food diet reduced excitability in MSNs from obesity-prone rats; this was mediated by an increase in I_*KIR*_. Thus, individual differences in obesity-susceptibility influence both basal excitability and how MSN excitability adapts to junk-food consumption.

### A. Basal differences in MSN excitability

Consistent with previous findings (Oginsky et al. 2016a) MSNs within the NAc core from obesity-prone rats fired more action potentials in response to current injection than MSNs from obesity-resistant rats. This was accompanied by a lower rheobase and greater input resistance in MSNs from obesity-prone vs. resistant rats. Furthermore, I_*A*_ amplitude was smaller in obesity-prone vs. obesity-resistant rats, consistent with increased excitability. Although it is unclear what is driving these basal differences, MSN excitability within the NAc core is comparable between obesity-prone and obesity-resistant during earlier stages of development (P30-40; Oginsky et al., 2016a). This suggests that there are not overt developmental differences between these groups, but rather that the emergence of differences during adulthood (P70-85) is more likely the result of complex gene x environment interactions. For example, in adulthood obesity-prone rats tend to be heavier and have more fat mass than obesity-resistant rats, even prior to overt obesity or metabolic dysregulation (Vollbrecht et al., 2015; Alonso-Caraballo et al., 2018). Though speculative, these differences in fat mass could influence NAc function via several systems and pathways including leptin-mediated modulation of dopamine systems (Leinninger et al. 2009; Leinninger et al. 2011; Opland et al. 2013; Perry et al. 2010). Of course, additional studies are needed to directly address this possibility.

### B. Effects of Junk-food diet on MSN excitability in obesity-prone and obesity-resistant rats

Ten days of junk-food consumption produced opposite effects on NAc core MSN excitability in obesity-prone and obesity-resistant rats compared to their chow fed counterparts. Specifically, junk-food increased MSN excitability in obesity-resistant groups (Figure 2). This was accompanied by a concomitant increase in input resistance, a reduction in I_*A*_ amplitude, and a shift in to the right of the I_*A*_ activation curve in MSNs from obesity-resistant rats given junk-food vs. chow (Figure 3). Thus, junk-food induced increase in excitability are due to a decrease in the activation of A-channels. In contrast, junk-food resulted in reduced MSN excitability in obesity-prone rats that was accompanied by increases in I_*KIR*_ (Figure 5), and no change in I_*A*_ compared to their chow fed counterparts. This is consistent with the observed increase in inward rectification and rheobase, and decrease in the RMP in obesity-prone JF vs chow groups (Figure 3). To our knowledge, only one similar study has been conducted. Fritz et al., (2018) gave mice free access to a western diet that was high in fat and sugar prior to examining intrinsic excitability in MSNs from the dorsal striatum (DS; measurements were not made from the NAc). They did not find any evidence for alterations in intrinsic excitability in the DS. Potential species differences aside, these data could suggest that diet-induced alterations vary by striatal sub-region, though caution should be taken as a number of differences between the studies (e.g., 16 weeks of western diet in Fritz et al., 2018, and 10 days here), make direct comparisons difficult.

#### Why might junk-food produce opposite effects on MSN intrinsic excitability in obesity-prone vs. obesity resistant rats?

Differences in baseline excitability described above may contribute to the opposing effects of junk-food in obesity-prone and obesity-resistant rats. For example, I_*A*_ amplitude is smaller in obesity-prone compared to obesity-resistant rats maintained on standard lab chow. Thus, further reductions in I_*A*_, and increases in excitability, may not be possible due to floor effects in obesity-prone rats. Instead, in obesity-prone rats junk-food produced a reduction in excitability by decreasing I_*KIR*_ magnitude. In contrast, basal I_*A*_ amplitude is larger in obesity-resistant rats, allowing for junk-food induced reductions in I_*A*_ to drive increases in intrinsic excitability. In this conceptualization, obesity-prone and obesity-resistant rats are beginning at two different points along the same inverted U shaped function of excitability, where obesity-prone rats begin near the peak and are pushed to the descending limb by junk-food consumption, whereas junk-food given to obesity-resistant rats pushes them up the ascending limb. This framework would suggest that sufficient junk-food consumption (and/or accompanying increases in adiposity) may also produce reductions in excitability in obesity-resistant rats. Unfortunately, it is difficult to induce substantial weight gain in obesity-resistant rats using this moderately fatty junk-food diet (~20%), and it is unknown whether or not high-fat diets that can produce substantive increases in fat mass in obesity-resistant rats are sufficient to alter intrinsic excitability (e.g., 60% high-fat; Alonso-Caraballo et al., 2019). None-the-less the current data demonstrate that interactions between predisposition and consumption of a junk-food diet produce distinct alterations in MSN excitability in obesity-prone vs. obesity-resistant males.

#### What could contribute to junk-food induced alterations in MSN intrinsic excitability?

Consumption of sugary and fatty foods enhances dopamine levels, leading to an increase in dopamine transmission to MSNs (Fordahl et al. 2016; Fritz et al. 2018) and enhanced sensitivity to psychostimulant-induced locomotor activity that relies in mesolimbic dopamine transmissions (Robinson et al. 2015). Dopamine receptor activation influences intrinsic excitability in part by regulating I_*KIR*_ and I_*A*_ (Azdad et al. 2009; Perez et al. 2006; Podda et al. 2010; Zhao et al. 2016). Thus, its likely that alterations in intrinsic excitability following junk-food consumption are secondary to dopamine receptor activation, although the specific mechanisms are likely complex (Hu et al. 2004; Nicola et al. 2000). Consistent with this idea, consumption of sugary, fatty foods also alters the expression and function of D2-type DA receptors in rodent models and human studies (Baladi et al. 2012; Robinson et al. 2015; Small 2009; Tomasi and Volkow 2013), though it should be noted that evidence for both increases and decreases (Huang et al. 2006; South and Huang 2008), as well as changes in D1-type DA receptor expression have been found (Alsi et al. 2010). Results here suggest that some of this heterogeneity of effects could be due to basal differences in MSN excitability. In addition, consumption of saturated fats alone are sufficient to alter mesolimbic systems and dopamine receptor function (Hryhorczuk et al. 2016). Thus, consumption of sugary fatty foods alone may be sufficient to alter mesolimbic systems. However, increases in fat mass and secondary effects of increased adiposity (such as insulin and leptin insensitivity) may also contribute.

While there is relatively little known about how natural rewards influence NAc intrinsic excitability, pharmacologically enhancing dopamine transmission (e.g., by repeated cocaine administration), also produces alterations in MSN intrinsic excitability. Specifically, repeated cocaine administration followed by a drug free period (withdrawal) results in a reduction in intrinsic excitability in the NAc (Dong et al. 2006; Hu et al. 2004). However, effects of cocaine vary in NAc core vs. shell subregions, and by duration of withdrawal (e.g., Mu et al., 2010). In addition cocaine-induced alterations in intrinsic excitability and have been linked to cocaine-seeking and craving that characterize addiction and to increases in AMPA-receptor mediated transmission that mediate these behaviors (Kourrich et al. 2015; Wolf 2010). Although food has no direct pharmacological actions in the brain, reductions in NAc intrinsic excitability and increases in AMPA receptor expression and function that are found in obesity-prone rats following junk-food consumption (Oginsky et al. 2016a) are generally similar to effects seen following cocaine. Of course, the degree to which similar vs. different circuits and neuronal sub-populations are altered by food vs. drug rewards remains to be determined (Ferrario, 2017).

### C. Summary and future directions

We show that eating junk-food alters MSN excitability by modulating voltage-gated potassium channels. Interestingly, this same diet manipulation produced opposite effects in obesity-resistant and obesity-prone rats. This heterogeneity may be related to basal difference in intrinsic excitability between these populations. In addition, future studies examining the role of obesity per se, as well as the persistence of these diet-induced alterations will further clarify the impact of obesogenic diets on mesolimbic function.

## Acknowledgments

This work was supported by NIDDK R01DK106188 and R01DK106188 to CRF; MFO was supported by NIDA T32DA007268 and NIDDK FDK112627A.

## Author Contributions

MFO designed conducted experiments, analyzed data, and wrote the manuscript. CRF designed experiments, analyzed data, and wrote the manuscript.

## REFERENCES

Alonso-Caraballo Y, Jorgensen ET, Brown T, and Ferrario CR. Functional and structural plasticity contributing to obesity: roles for sex, diet, and individual susceptibility. Curr Opin Behav Sci 23: 160–170, 2018.

Alsi J, Olszewski PK, Norbck AH, Gunnarsson ZE, Levine AS, Pickering C, and Schith HB. Dopamine D1 receptor gene expression decreases in the nucleus accumbens upon long-term exposure to palatable food and differs depending on diet-induced obesity phenotype in rats. Neuroscience 171: 779–787, 2010.

Azdad K, Chvez M, Don Bischop P, Wetzelaer P, Marescau B, De Deyn PP, Gall D, and Schiffmann SN. Homeostatic plasticity of striatal neurons intrinsic excitability following dopamine depletion. PLoS One 4: e6908, 2009.

Baladi MG, Daws LC, and France CP. You are what you eat: influence of type and amount of food consumed on central dopamine systems and the behavioral effects of direct-and indirect-acting dopamine receptor agonists. Neuropharmacology 63: 76–86, 2012.

Bassareo V, Cucca F, Musio P, Lecca D, Frau R, and Di Chiara G. Nucleus accumbens shell and core dopamine responsiveness to sucrose in rats: role of response contingency and discriminative/conditioned cues. Eur J Neurosci 41: 802–809, 2015.

Demos KE, Heatherton TF, and Kelley WM. Individual differences in nucleus accumbens activity to food and sexual images predict weight gain and sexual behavior. J Neurosci 32: 5549–5552, 2012.

Derman RC, and Ferrario CR. Enhanced incentive motivation in obesity-prone rats is mediated by NAc core CP-AMPARs. Neuropharmacology 131: 326–336, 2018a.

Derman RC, and Ferrario CR. Junk-food enhances conditioned food cup approach to a previously established food cue, but does not alter cue potentiated feeding; implications for the effects of palatable diets on incentive motivation. Physiol Behav 192: 145–157, 2018b.

Dong Y, Green T, Saal D, Marie H, Neve R, Nestler EJ, and Malenka RC. CREB modulates excitability of nucleus accumbens neurons. Nat Neurosci 9: 475–477, 2006.

Ferrario CR. Food Addiction and Obesity. Neuropsychopharmacology 42: 361, 2017.

Fordahl SC, Locke JL, and Jones SR. High fat diet augments amphetamine sensitization in mice: Role of feeding pattern, obesity, and dopamine terminal changes. Neuropharmacology 109: 170–182, 2016.

Fritz BM, Muoz B, Yin F, Bauchle C, and Atwood BK. A High-fat, High-sugar ‘Western’ Diet Alters Dorsal Striatal Glutamate, Opioid, and Dopamine Transmission in Mice. Neuroscience 372: 1–15, 2018.

Henry DJ, and White FJ. Electrophysiological correlates of psychomotor stimulant-induced sensitization. Ann N Y Acad Sci 654: 88–100, 1992.

Hryhorczuk C, Florea M, Rodaros D, Poirier I, Daneault C, Des Rosiers C, Arvanitogiannis A, Alquier T, and Fulton S. Dampened Mesolimbic Dopamine Function and Signaling by Saturated but not Monounsaturated Dietary Lipids. Neuropsychopharmacology 41: 811–821, 2016.

Hu XT, Basu S, and White FJ. Repeated cocaine administration suppresses HVA-Ca2+ potentials and enhances activity of K+ channels in rat nucleus accumbens neurons. J Neurophysiol 92: 1597–1607, 2004.

Huang XF, Zavitsanou K, Huang X, Yu Y, Wang H, Chen F, Lawrence AJ, and Deng C. Dopamine transporter and D2 receptor binding densities in mice prone or resistant to chronic high fat diet-induced obesity. Behav Brain Res 175: 415–419, 2006.

Jin X, Cui N, Zhong W, Jin XT, and Jiang C. GABAergic synaptic inputs of locus coeruleus neurons in wild-type and Mecp2-null mice. Am J Physiol Cell Physiol 304: C844–857, 2013.

John J, and Manchanda R. Modulation of synaptic potentials and cell excitability by dendritic KIR and KAs channels in nucleus accumbens medium spiny neurons: a computational study. J Biosci 36: 309–328, 2011.

Kourrich S, Calu DJ, and Bonci A. Intrinsic plasticity: an emerging player in addiction. Nat Rev Neurosci 16: 173–184, 2015.

Kourrich S, and Thomas MJ. Similar neurons, opposite adaptations: psychostimulant experience differentially alters firing properties in accumbens core versus shell. J Neurosci 29: 12275–12283, 2009.

Leinninger GM, Jo YH, Leshan RL, Louis GW, Yang H, Barrera JG, Wilson H, Opland DM, Faouzi MA, Gong Y, Jones JC, Rhodes CJ, Chua S, Diano S, Horvath TL, Seeley RJ, Becker JB, Mnzberg H, and Myers MG. Leptin acts via leptin receptor-expressing lateral hypothalamic neurons to modulate the mesolimbic dopamine system and suppress feeding. Cell Metab 10: 89–98, 2009.

Leinninger GM, Opland DM, Jo YH, Faouzi M, Christensen L, Cappellucci LA, Rhodes CJ, Gnegy ME, Becker JB, Pothos EN, Seasholtz AF, Thompson RC, and Myers MG. Leptin action via neurotensin neurons controls orexin, the mesolimbic dopamine system and energy balance. Cell Metab 14: 313–323, 2011.

Levin BE, Dunn-Meynell AA, Balkan B, and Keesey RE. Selective breeding for diet-induced obesity and resistance in Sprague-Dawley rats. Am J Physiol 273: R725–730, 1997.

McCutcheon JE, and Roitman MF. Mode of Sucrose Delivery Alters Reward-Related Phasic Dopamine Signals in Nucleus Accumbens. ACS Chem Neurosci 10: 1900–1907, 2019.

Mermelstein PG, Song WJ, Tkatch T, Yan Z, and Surmeier DJ. Inwardly rectifying potassium (IRK) currents are correlated with IRK subunit expression in rat nucleus accumbens medium spiny neurons. J Neurosci 18: 6650–6661, 1998.

Mu P, Moyer JT, Ishikawa M, Zhang Y, Panksepp J, Sorg BA, Schlter OM, and Dong Y. Exposure to cocaine dynamically regulates the intrinsic membrane excitability of nucleus accumbens neurons. J Neurosci 30: 3689–3699, 2010.

Nicola SM, Surmeier J, and Malenka RC. Dopaminergic modulation of neuronal excitability in the striatum and nucleus accumbens. Annu Rev Neurosci 23: 185–215, 2000.

Nisenbaum ES, and Wilson CJ. Potassium currents responsible for inward and outward rectification in rat neostriatal spiny projection neurons. J Neurosci 15: 4449–4463, 1995.

Oginsky MF, Goforth PB, Nobile CW, Lopez-Santiago LF, and Ferrario CR. Eating ‘Junk-Food’ Produces Rapid and Long-Lasting Increases in NAc CP-AMPA Receptors: Implications for Enhanced Cue-Induced Motivation and Food Addiction. Neuropsychopharmacology 41: 2977–2986, 2016a.

Oginsky MF, Maust JD, Corthell JT, and Ferrario CR. Enhanced cocaine-induced locomotor sensitization and intrinsic excitability of NAc medium spiny neurons in adult but not in adolescent rats susceptible to diet-induced obesity. Psychopharmacology (Berl) 233: 773–784, 2016b.

Opland D, Sutton A, Woodworth H, Brown J, Bugescu R, Garcia A, Christensen L, Rhodes C, Myers M, and Leinninger G. Loss of neurotensin receptor-1 disrupts the control of the mesolimbic dopamine system by leptin and promotes hedonic feeding and obesity. Mol Metab 2: 423–434, 2013.

Perez MF, White FJ, and Hu XT. Dopamine D(2) receptor modulation of K(+) channel activity regulates excitability of nucleus accumbens neurons at different membrane potentials. J Neurophysiol 96: 2217–2228, 2006.

Perry ML, Leinninger GM, Chen R, Luderman KD, Yang H, Gnegy ME, Myers MG, and Kennedy RT. Leptin promotes dopamine transporter and tyrosine hydroxylase activity in the nucleus accumbens of Sprague-Dawley rats. J Neurochem 114: 666–674, 2010.

Podda MV, Riccardi E, D’Ascenzo M, Azzena GB, and Grassi C. Dopamine D1-like receptor activation depolarizes medium spiny neurons of the mouse nucleus accumbens by inhibiting inwardly rectifying K+ currents through a cAMP-dependent protein kinase A-independent mechanism. Neuroscience 167: 678–690, 2010.

Robinson MJ, Burghardt PR, Patterson CM, Nobile CW, Akil H, Watson SJ, Berridge KC, and Ferrario CR. Individual Differences in Cue-Induced Motivation and Striatal Systems in Rats Susceptible to Diet-Induced Obesity. Neuropsychopharmacology 40: 2113–2123, 2015.

Sekerli M, Del Negro CA, Lee RH, and Butera RJ. Estimating action potential thresholds from neuronal time-series: new metrics and evaluation of methodologies. IEEE Trans Biomed Eng 51: 1665–1672, 2004.

Small DM. Individual differences in the neurophysiology of reward and the obesity epidemic. Int J Obes (Lond) 33 Suppl 2: S44–48, 2009.

South T, and Huang XF. High-fat diet exposure increases dopamine D2 receptor and decreases dopamine transporter receptor binding density in the nucleus accumbens and caudate putamen of mice. Neurochem Res 33: 598–605, 2008.

Stice E, Spoor S, Bohon C, Veldhuizen MG, and Small DM. Relation of reward from food intake and anticipated food intake to obesity: a functional magnetic resonance imaging study. J Abnorm Psychol 117: 924–935, 2008.

Tomasi D, and Volkow ND. Striatocortical pathway dysfunction in addiction and obesity: differences and similarities. Crit Rev Biochem Mol Biol 48: 1–19, 2013.

Uchimura N, Cherubini E, and North RA. Inward rectification in rat nucleus accumbens neurons. J Neurophysiol 62: 1280–1286, 1989.

Uchimura N, and North RA. Muscarine reduces inwardly rectifying potassium conductance in rat nucleus accumbens neurones. J Physiol 422: 369–380, 1990.

Vainik U, Dagher A, Dub L, and Fellows LK. Neurobehavioural correlates of body mass index and eating behaviours in adults: a systematic review. Neurosci Biobehav Rev 37: 279–299, 2013.

Vollbrecht PJ, Nobile CW, Chadderdon AM, Jutkiewicz EM, and Ferrario CR. Pre-existing differences in motivation for food and sensitivity to cocaine-induced locomotion in obesity-prone rats. Physiol Behav 152: 151–160, 2015.

Wilson CJ, and Kawaguchi Y. The origins of two-state spontaneous membrane potential fluctuations of neostriatal spiny neurons. J Neurosci 16: 2397–2410, 1996.

Wolf ME. The Bermuda Triangle of cocaine-induced neuroadaptations. Trends Neurosci 33: 391–398, 2010.

Yokum S, Ng J, and Stice E. Attentional bias to food images associated with elevated weight and future weight gain: an fMRI study. Obesity (Silver Spring) 19: 1775–1783, 2011.

Zhao B, Zhu J, Dai D, Xing J, He J, Fu Z, Zhang L, Li Z, and Wang W. Differential dopaminergic regulation of inwardly rectifying potassium channel mediated subthreshold dynamics in striatal medium spiny neurons. Neuropharmacology 107: 396–410, 2016.

